# Road density simplifies regional food webs

**DOI:** 10.1101/2024.03.07.583843

**Authors:** Frederico Mestre, Vinicius A. G. Bastazini, Fernando Ascensao

## Abstract

Roads stands as major threats to biodiversity, affecting the functioning of ecosystems and their provision of ecosystem services. Understanding how road-related impacts affect the dynamics of ecological interactions is essential to help manage human impacts on biodiversity, but such studies remain largely unexplored. We investigated the intricate relationship between roads, species vulnerability to road density, and the ensuing effects on food webs across Europe. Utilizing species-specific road density thresholds and trophic interaction data, we constructed regional food webs to assess the potential loss of trophic interactions due to roadkill. Our analysis, encompassing 551 species across three trophic levels, revealed spatially varied impacts, with areas surrounding major cities facing drastic losses of trophic interaction exceeding 90%. Notably, 191 species may be affected by trophic interaction losses (loss of prey or predator). Apex predators exhibited lower direct impacts from road density, while basal-level species seem to be more exposed to direct road-related effects, potentially triggering a cascade of interaction disruptions. Our findings emphasize the need for informed road infrastructure development and targeted conservation strategies to mitigate the negative impacts of roads and traffic, thereby preserving the integrity of ecological networks. This research offers valuable insights for policymakers and conservation practitioners globally by identifying critical areas where road-induced cascade effects may be most pronounced and highlighting trophic groups of species that may be at higher risk.

## Introduction

Roads can play a vital role in human development by allowing access to important services, such as education and health care, further contributing to economic growth by enabling trade (Gachassin et al., 2010; Iimi et al., 2015; Sewell et al., 2019). However, roads and traffic stand as a major threat to biodiversity, leading to habitat fragmentation and degradation (Forman and Alexander, 1998; Maxwell et al., 2016; Van der Ree et al., 2015). Notably, the expansion of road networks and the associated increase in traffic drastically affects wildlife populations, primarily due to animal-vehicle collisions (Grilo et al., 2020; Morelli et al., 2020). In developed regions like Europe, where the density of roads rank among the highest globally (Meijer et al., 2018), roads exert a considerable impact on wildlife. For example, recent estimates point to over 200 million individual birds and mammals being road-killed on European paved roads every year (Grilo et al., 2020).

Road-related impacts are likely to ripple through regional ecological communities when species’ interactions are weakened or even lost following reductions in population abundance or local extinctions (Barrientos et al., 2020). A simplification of ecological network structure such as food webs (Dunne et al., 2002; Mestre et al., 2022), can change fluxes of matter and energy (Barnes et al., 2018), affecting the functioning of ecosystems, ultimately disrupting the provision of ecosystem services (Barbosa et al., 2017; Tylianakis et al., 2008). This simplification of trophic chains can further impact species indirectly by altering prey density or reducing predator pressure (Leighton et al., 2023). Therefore, understanding how roads and traffic affect the dynamics of ecological interactions, particularly food webs, is crucial for managing human impacts on biodiversity, especially those associated with transport infrastructure. However, the study of road-induced impacts on ecological networks remains largely unexplored, likely due to the logistical and financial complexities of monitoring numerous regional food webs across varying road density scenarios and over large spatial extents.

Previous research established the generalized theoretical population model for deriving species-specific vulnerability to roads based on road density thresholds, beyond which local populations may not persist (Borda-de-Água et al., 2011). This theoretical framework was recently used to calculate thresholds of road density for a collection of European birds and mammals (Grilo et al., 2020) (see Methods for further details). Some examples of the thresholds of road density include 0.91 km/km^2^ for Iberian lynx (*Lynx pardinus*), 1.43 km/km^2^ for golden eagle (*Aquila chrysaetos*), 2.51 km/km^2^ for wild cat (*Felis silvestris*), and 3.00 km/km^2^ for the wild rabbit (*Oryctolagus cuniculus*) (all species-specific threshold values are available in Grilo et al. (2020)). We used these estimated thresholds to infer where potential losses in species’ occurrences may occur and, consequently, the interactions they were involved in (Fig. 1).

**Figure 1.**
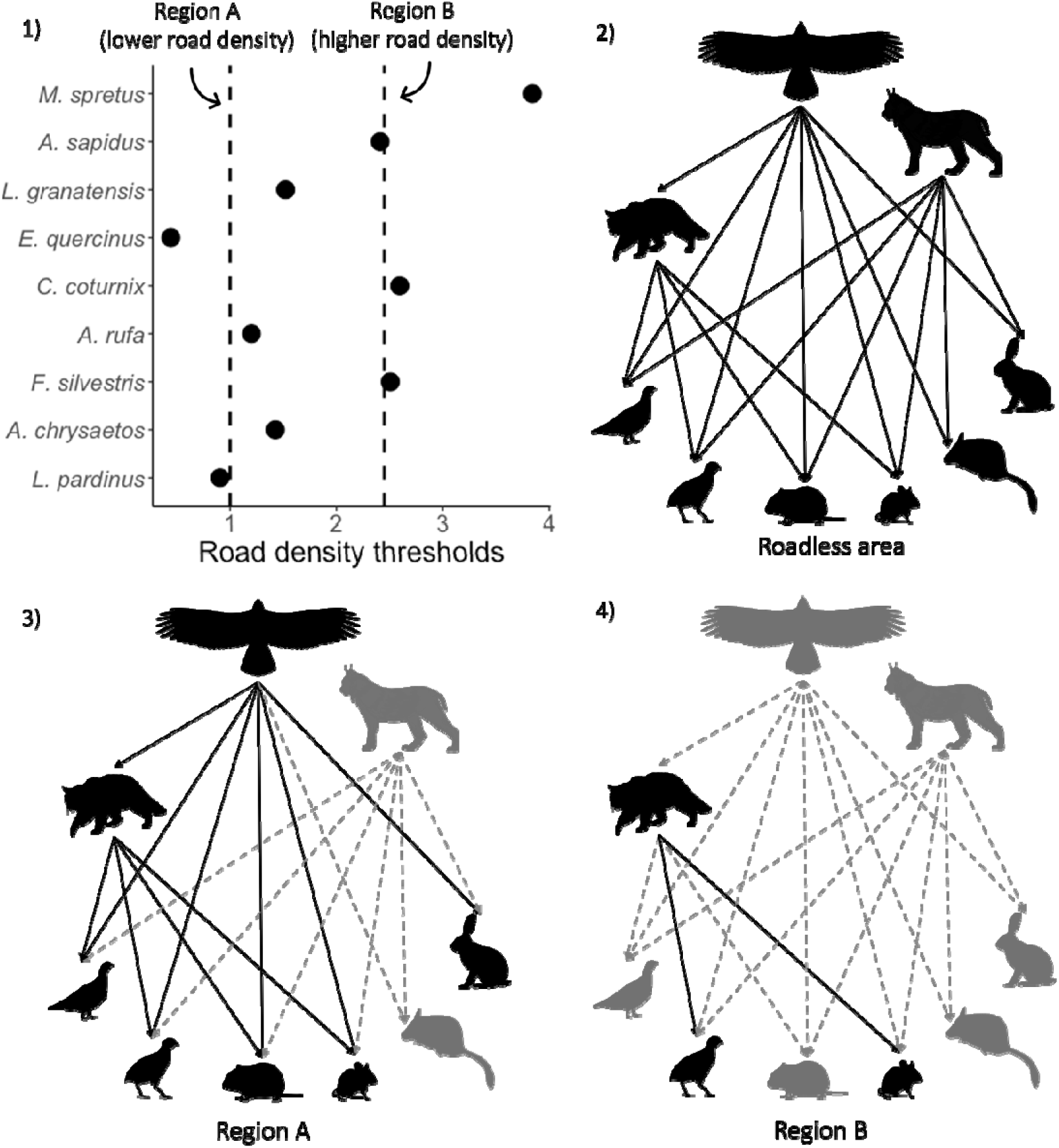
Road-induced simplification of food webs. The diagram illustrates the potential loss of interactions resulting from road density exceeding species-specific thresholds. In Panel 1, the road density thresholds for nine species are presented. The garden dormouse (*E. quercinus*) shows the lowest estimated road density threshold (0.4 km/km^2^), while the Algerian mouse (*M. spretus*) has the highest threshold (3.8 km/km^2^). These species participate in the food web depicted in Panel 2 (network not exhaustive), representing a scenario without roads. For two regions with lower and higher road densities (Panels 3 and 4; also indicated in Panel 1), the network may simplify: some species face a risk of local extinction (shown in grey) when road densities exceed their thresholds, increasing the likelihood of roadkill and threatening their survival. This leads to a loss of interactions (indicated by dashed lines). In contrast, species depicted in black are more likely to persist, as their road density thresholds exceed the levels present in the region. In the example provided, the Iberian lynx (*L. pardinus*) and garden dormouse may face extinction in region A. In region B, golden eagle (*A. chrysaetos*) and its prey, including red-legged partridge (*A. rufa*), southern water vole (*A. sapidus*), and Iberian hare (*L. granatensis*), could also be at risk, disrupting predator-prey dynamics. Wild cat (*F. silvestris*), common quail (*C. coturnix*), and Algerian mouse might persist even at the highest road densities, as can their trophic interactions. Silhouettes from phylopic (www.phylopic.org).

Using the combined information on road density thresholds and food webs, we investigated how food webs can be imperiled when road density surpasses the species-specific thresholds beyond which local populations may not persist. Specifically, we sought to evaluate the extent of potential interaction loss across Europe and trophic levels; and compare, for each species, the amount of distribution area that could be lost or degraded due to road density exceeding *i*) its specific vulnerability threshold (direct effect) and *ii*) that of the species that interact with it i.e., prey and/or predators (indirect effect). We expected a higher loss of interactions and higher degradation of range area with increasing road density values. However, given the historical presence of roads in Europe and the higher vulnerability of apex predators to roadkill effects (Rytwinski and Fahrig, 2012), we predicted a higher direct effect for lower trophic levels and a higher vulnerability of interactions at higher trophic levels due to indirect effects.

## Methods

### Species vulnerability to roads

Species-specific thresholds for road density were obtained for birds and mammals (Grilo et al., 2020). The authors estimated long-term vulnerability to roadkill using the theoretical demographic model developed by Borda-de-Água et al. (2011), which differentiates between landscapes with roads and non-roads:

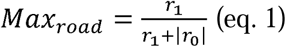

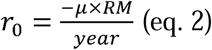

Following Grilo et al. (2020), *r*_1_ is the intrinsic population growth rate and *r*_0_ is the annual rate of population decay on paved roads, where µ is the species’ annual natural mortality rate given its longevity, and *RM* is a scaling factor for road mortality assuming that roads are unsuitable environments for species to persist, with *RM* = 1000 having been the value used, in accordance to previously assumed by (Ceia-Hasse et al., 2017). According to the authors, in this model, non-road areas support positive population growth rates, while road areas lead to rapid population declines, assuming roads are unsuitable habitats. The intrinsic population growth rate (*r*_1_) is calculated using a simplified version of the Euler equation that incorporates species traits such as maturity age and offspring, as well as the annual natural mortality rate, based on species longevity (Borda-de-Água et al., 2011; Grilo et al., 2020). Using this approach, the authors calculated species-specific maximum road density thresholds (eq. 1), beyond which population persistence is unlikely. Importantly, both the authors and our study consider only paved roads, as these are where roadkill is more likely to occur due to generally higher traffic volumes [see Borda-de-Água et al. (2011) for further details on the modelling procedure, and Grilo et al. (2020) for further details on the estimation of species-specific thresholds]. We regarded the road density threshold as an indicator that enables a species-specific ranking of regions where species are unlikely to persist, potentially leading to the loss of their ecological interactions.

### Building local potential food webs and risk of interaction disruption

We built local trophic structures using a grid of 50 x 50 km cells overlaid upon the European region (n=4,743 cells), the same used by Grilo et al. (2020). Building local potential food webs was guided by the observed interactions outlined in the TETRA-EU metaweb (Maiorano et al., 2020). This metaweb is based on observed interactions obtained from the scientific literature and summarizes the potential trophic interactions for tetrapod species (mammals, breeding birds, reptiles, and amphibians) across Europe and the Northern Mediterranean basin. If a trophic interaction between two species was documented in the metaweb, it was considered indicative of their potential interaction within the regional network (i.e., at the grid cell level). We characterized the trophic level of each species in the metaweb using a categorical classification, dividing species into three groups: top-level (species that only have prey), intermediate level (species with both predators and prey), and basal (species that only have predators). We did this by resorting to the functions ‘BasalNodes’, ‘IntermediateNodes’, and ‘TopLevelNodes’ of the cheddar R package (Hudson et al., 2013).

We then used the road density thresholds to predict potential points of trophic interaction disruption in the event of species extinction or severe depletion. For each species, we assumed that grid cells exceeding their species-specific maximum road density threshold, i.e., beyond which long-term population persistence becomes improbable (Borda-de-Água et al., 2011), would exhibit an increased probability of losing trophic interactions associated with that species. Extinction simulation was conducted using the function ‘RemoveNodes’ of the cheddar R package (Hudson et al., 2013).

All the computations were conducted in R (R Core Team, 2021), using the R packages cheddar (for network analysis) (Hudson et al., 2013), igraph (for network analysis) (Csárdi et al., 2023), taxize (to combine both datasets, the vulnerability dataset and the metaweb) (Chamberlain and Szöcs, 2013), and terra (for R based spatial analysis and for creating the output shapefiles) (Hijmans, 2023). Images were produced in QGIS (2020) and R (R Development Core Team, 2022). The dataset is available in <u>FIGSHARE</u>. All code is available on <u>GitHub</u>.

## Results

### General patterns

Our pool of species, for which a threshold of vulnerability to road density was available and referenced in the TETRA-EU metaweb, consisted of 551 species (349 birds and 202 mammals). This pool comprised 12 top-level species (2.2% of the total), 107 intermediate-level species (19.4%), and 432 basal-level species (78.4%) (see Supplementary material S1 for species names). The three trophic levels currently exhibit varying distributions across Europe (Fig. 2A-C), notably with higher species richness in regions characterized by lower road density (Fig. 2D), particularly top-level species. This pool of species assembled altogether a mean (±SD) of 2,057±819 predator-prey interactions per grid cell across Europe. A greater number of interactions were observed in mountainous regions like the Pyrenees, Alps, and Carpathians, alongside areas in eastern Europe, regions that exhibit lower road density (Fig. 2E).

**Figure 2.**
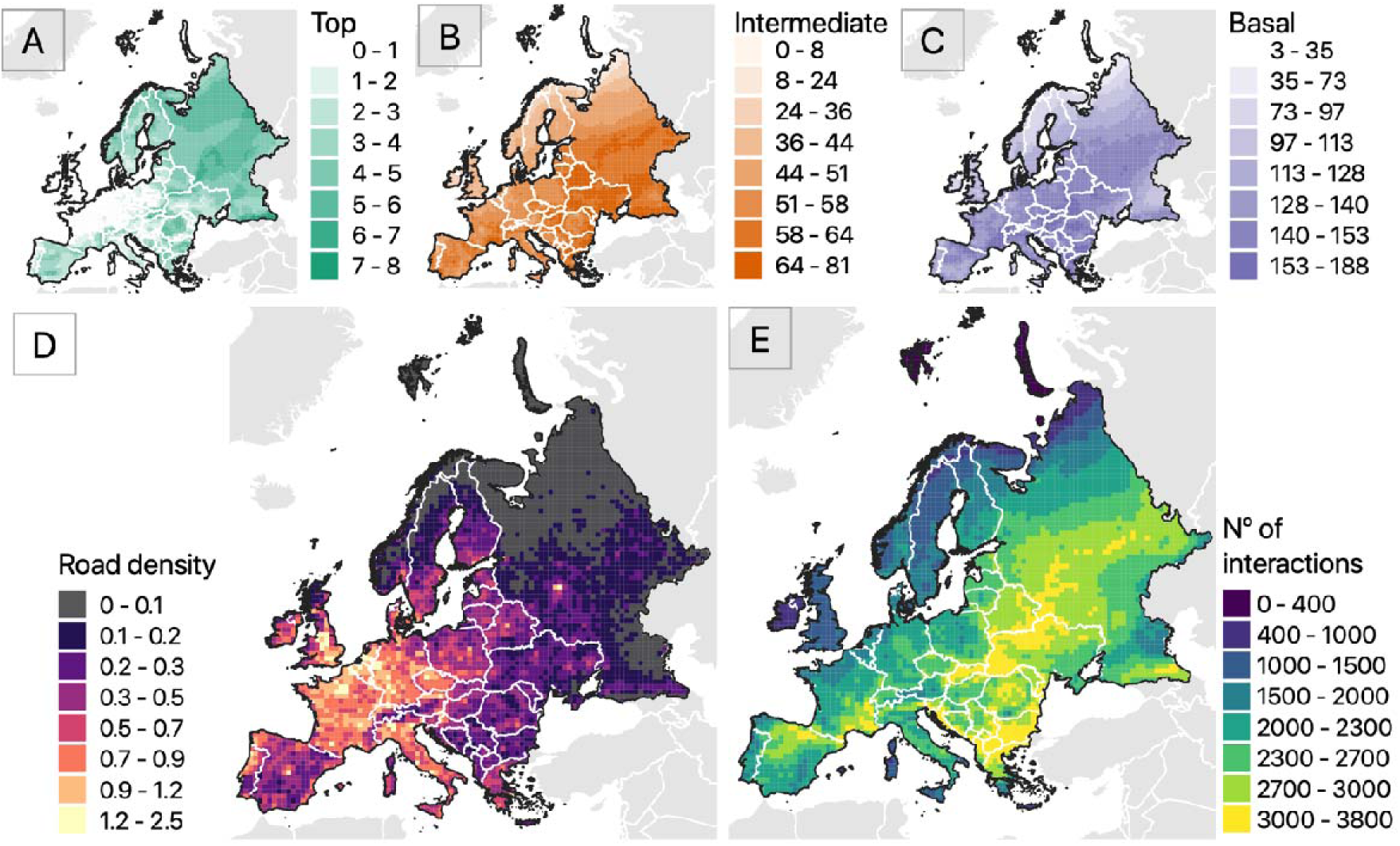
Richness of top, intermediate and basal level species (A-C), road density in km/km^2^ (D), and estimated number of interactions (E) per grid cell, across Europe. All maps use a grid cell resolution of 50×50 km. White lines depict country borders.

### Trophic interaction loss

On average, 4.5% (±9.4) of interactions in our regional food webs face potential loss due to road impacts i.e., when road density surpasses the species-specific vulnerability threshold of either the prey or predator engaged in the interaction. However, the simplification of food webs is likely to vary spatially, being particularly intense in areas surrounding large European cities. In these areas, road density can reach 2.5 km/km² (see Fig. 2D), resulting in the loss of over 90% of potential interactions (Fig. 3A). The loss of interactions in the local food webs seems to have a strong relationship with road density, reaching a 30% loss when road density approaches 1 km/km² (Fig. 3B).

**Figure 3.**
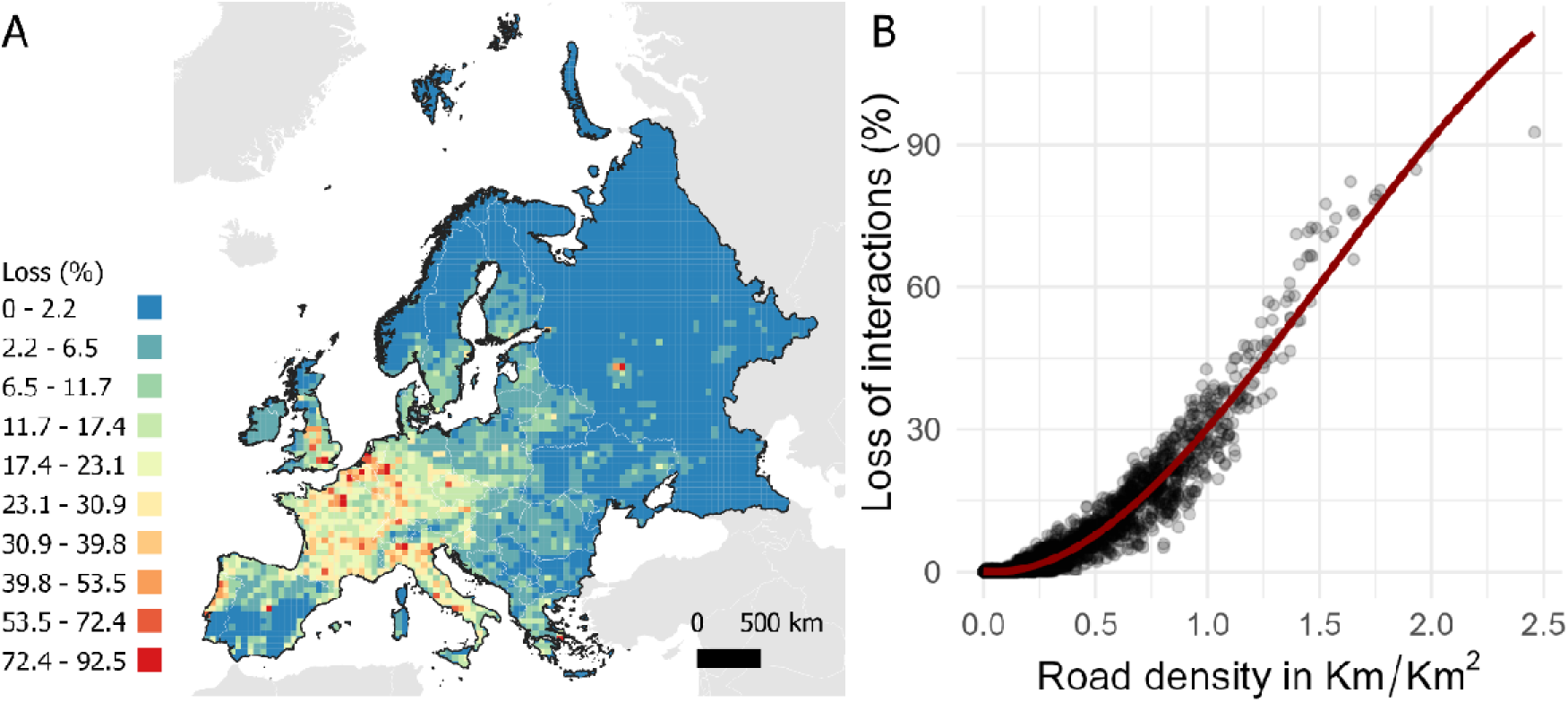
A) Expected loss of interactions due to road impacts; B) Relation between road density and interaction loss (red line depicts the smooth line using a polynomial spline, here used for better visualization of the relation between both variables).

### Effects within species range

Among the species examined, 81 (15% of the total) had over 10% of their range with road density surpassing their vulnerability threshold, putting them at significant risk of local extinction due to direct road impacts. These include one top-level species, 12 intermediate species, and 68 basal species (Fig. 4). Notably, the apex predator booted eagle *(Hieraaetus pennatus*) may disappear from 16% of its current range (see Supplementary material S1). The intermediate-level species short-toed snake eagle *(Circaetus gallicus)* and garden dormouse *(Eliomys quercinus)* may experience a substantial loss or degradation within over one-third of their range. For the basal-level species hazel grouse *(Tetrastes bonasia)* and both mole rats *(Spalax graecus* and *S. zemni)*, road densities surpass their vulnerability threshold in nearly half the range (grouse) or the full range (mole rats), leading to potential local extinctions across their entire territories.

**Figure 4.**
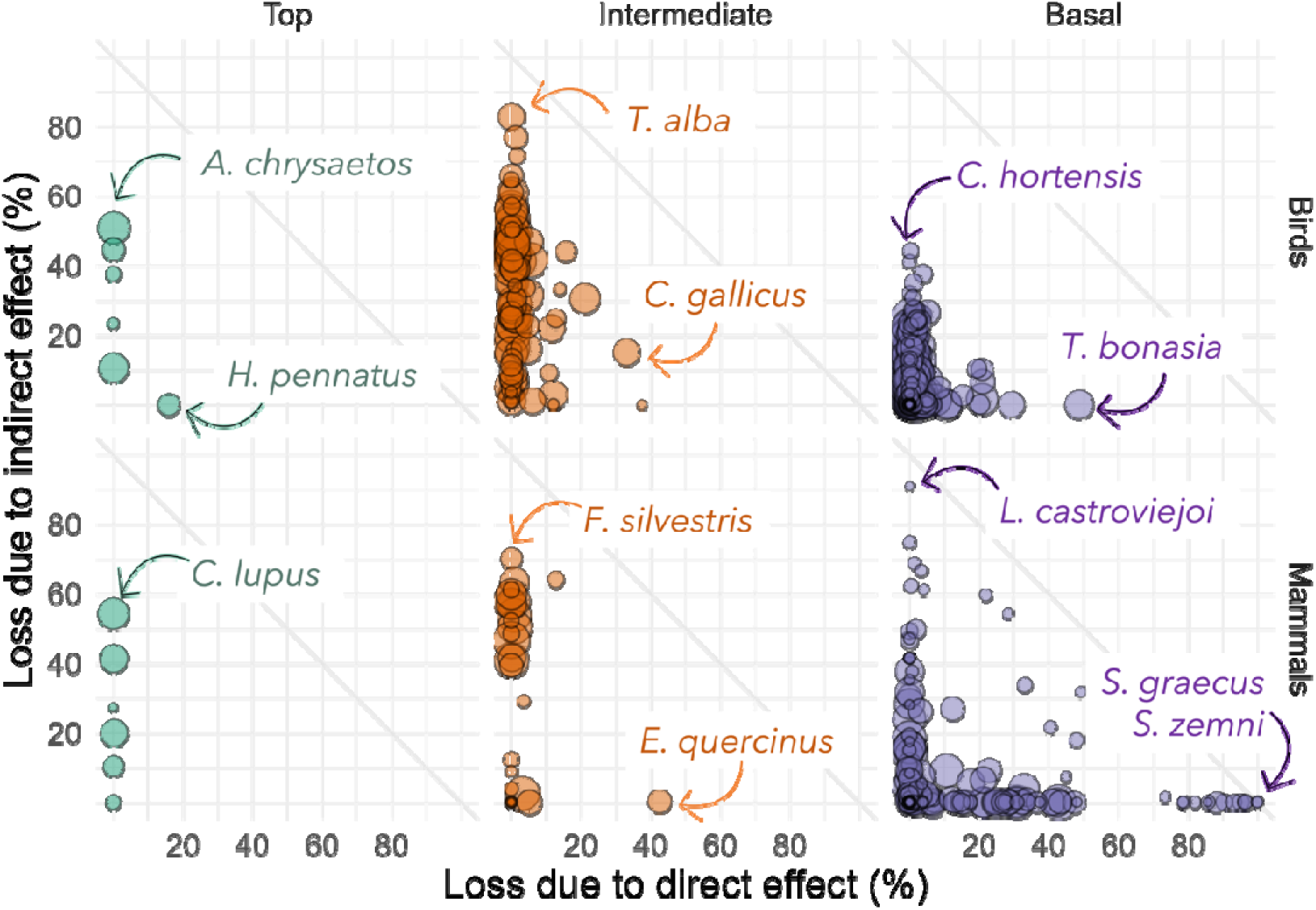
Range area with potential trophic interaction losses due to direct and indirect effects of roads, by trophic level (top, intermediate and basal). The circle radius is proportional to the range of the species. Grey diagonal line represents 100% of range areas affected by combination of direct and indirect effects. Highlighted species are mentioned in the text.

On the other hand, 191 species (35% of the total), including 10 top-level, 81 intermediate-level, and 100 basal-level species, exhibit more than 10% of their range at risk of losing trophic interactions due to indirect effects (Fig. 4). Notably, two top-level species, the golden eagle and the wolf (*Canis lupus*), may experience interaction losses with prey in over half their territory (Fig. 4). For example, the golden eagle is mostly affected by potential interaction losses with the hazel grouse (*Bonasa bonasia*; 16% of the total interactions lost), the European snow vole (*Chionomys nivalis*; 8%), the European mole (*Talpa europaea*; 7%) and the garden dormouse (7%). As for the wolf, it will potentially lose interactions mostly with hazel grouse (13% of total interactions), Eurasian pygmy shrew (*Sorex minutus*; 13%) and the common shrew (*Sorex araneus*; 8%).

Likewise, intermediate-level species, such as barn owl (*Tyto alba*) and wild cat (*Felis silvestris*), may experience losses of trophic interactions in over 70% of their range. The barn owl may lose most of its interactions as a consumer, namely with the Eurasian pygmy shrew (8%), the whiskered bat (*Myotis mystacinus*; 5%) and the European mole (5%). Conversely, as resource, the barn owl may lose interactions with the Eurasian goshawk (*Accipiter gentilis*; 90%) and the tawny owl (*Strix aluco*; 10%). As for the wild cat, it will lose most of its interactions as a predator with the pygmy shrew (14%), the common shrew (8%) and the hazel grouse (8%). As a resource, the wild cat will lose interactions with the Eurasian lynx (*Lynx lynx*; 75%) and the golden eagle (25%). Finally, the basal-level species, Western Orphean warbler *(Curruca hortensis)* and broom hare (*Lepus castroviejoi*), may lose interactions as prey in 44% and 91% of their range, respectively (Fig. 4). The Western Orphean warbler may experience interaction losses mostly from Eurasian scops owl (*Otus scops*; 39%), booted eagle (26%), woodchat shrike (*Lanius senator*; 17%) or Montagu’s Harrier (*Circus pygargus*; 7%). On the other hand, the broom hare experiences interaction losses exclusively with the short-toed snake eagle.

These findings strongly imply that the detrimental impacts of road density extend far beyond what meets the eye. For instance, top-level species currently face a minor risk of depletion due to roadkill. However, they may suffer from indirect effects across a significant portion of their range, where at least one prey species goes extinct due to roadkill (Fig. 4).

## Discussion

Road-related effects are expected to have a direct impact on species occurrence and abundance (Andrasi et al., 2021; Ascensão et al., 2022; Borda-de-Água et al., 2011; Forman and Alexander, 1998). Roadkill, in particular, can be a major driver of population depletion and reduce the population persistence in road surrounding areas (Ascensão and Desbiez, 2022; Barrientos et al., 2021; Moore et al., 2023). Yet, beyond such population-level effects, its repercussions are likely to also exert profound consequences on ecological networks. Our results indicate that increasing road density may significantly reduce the number of trophic interactions. This simplification of the food web is due not only to the direct depletion of populations but also to the indirect effects on food chains, such as the elimination of predators or prey. Furthermore, this loss of interactions occurs differently across space, with a greater incidence near large cities, and areas coinciding with higher road density, such as across Belgium, France, Germany, Italy, and the north of the Iberian Peninsula. Such imbalanced losses may have far-reaching implications for regional food webs, which could ultimately culminate in local extinctions, disturbing ecosystem functioning, and disrupting the delivery of essential ecosystem services.

Overall, top and intermediate species seem to follow a pattern dominated by loss of interactions due to the indirect effects of roads, with intermediate species having higher range area affected by losses than top-level species. Apart from the booted eagle, which presently inhabits an extensive region marked by road densities exceeding its estimated vulnerability threshold, the other top-level species like the golden eagle, brown bear (*Ursus arctos*), or Eurasian lynx, face minimal risk of being depleted due to direct road-related effects, as their current ranges have low road densities. This implies that impacts such as roadkill may not directly affect the persistence of the majority of top-level species. This is not surprising given that such apex predators are already restricted to habitats with relatively lower human impact and activity, and consequently with low road density areas (Bleyhl et al., 2021; Cretois et al., 2021). Conversely, basal species seem to be highly impacted by direct effects, therefore facing local extinction risk, and potentially eliminating the interactions with their predators. Yet, several basal species may also benefit from a release of predator interactions in their range areas.

Research has demonstrated that habitat fragmentation, in general, affects mostly consumer species, with larger body sizes, at higher trophic levels (Rytwinski and Fahrig, 2013, 2012). These species will generally go extinct faster than species with smaller body sizes and at lower trophic levels, even if having better dispersal ability (Liao et al., 2017; Ryser et al., 2019). On the other hand, despite the widespread suitable habitat for large carnivores, such as the brown bear and the Iberian lynx in Iberian Peninsula, human pressure is limiting the potential range expansion of these species (Pratzer et al., 2023). Likewise, wolves and other large carnivores are known to avoid sources of human disturbance, including high-road-density areas (Bleyhl et al., 2021; Cretois et al., 2021; Rio-Maior et al., 2019). This might explain why in our dataset top-level species are mostly restricted to lower road density areas.

Despite the long debate on the role of bottom-up versus top-down control in food webs (Leroux and Loreau, 2015), some evidence suggests that human activities tend to amplify bottom-up effects while weakening top-down trophic cascades in terrestrial ecosystems (Muhly et al., 2013). Our results seem to corroborate this notion, as they indicate that the impact of roads is particularly strong on prey species, disrupting their populations, thereby affecting higher trophic levels in the regional networks. Similar to our findings, previous research has also shown that prey species can benefit from lower exposition to predation pressure (Liao et al., 2017), and that habitat fragmentation – and probably also high road density areas – leads to ecological network simplification (Pires et al., 2023). However, the impact of roads on intermediate and basal-level generalist species may have stronger implication for food web dynamics, as generalist species play important roles in structuring the topology of ecological networks (Bastazini et al., 2019; Gaston and Fuller, 2008; Memmott et al., 2004; Poisot et al., 2013; Richmond et al., 2005; Valiente-Banuet et al., 2015). These species interact with more species than specialist species, thereby creating a more cohesive network (Bastazini et al., 2019; Martín González et al., 2010). By linking otherwise disconnected sub-networks, generalists enhance the overall connectivity, which is likely to increase their ability to respond to perturbations, increasing their stability (Bascompte et al., 2003).

We acknowledge that there are three limitations in our study. Firstly, although the metaweb approach offers valuable insights into species’ ecological associations, it overlooks the spatial and ecological context of these interactions (Chamberlain et al., 2014; Liu and Gaines, 2022). The strength and distribution of species interactions may differ across different populations and habitats due to environmental conditions and the presence of other species. For example, the Iberian lynx may have a few prey species (27 prey species according to metaweb used). However, its persistence is particularly dependent on the presence of one key prey, the wild rabbit, which constitutes 75%–99% of its diet (Fedriani et al., 1999). As such, losing interactions with other prey would probably not have relevant consequences for the persistence for this top predator. Incorporating interaction strength into food webs, by using weighted networks instead of unweighted ones, would improve the realism of our models. However, there is no available information that would allow us to construct such networks.

On the other hand, the database on trophic interactions builds on species’ co-occurrence. As so, we do not fully account for the spatial structure of these biotic interactions (e.g., Poisot et al., 2015). The strength of interactions between two species can vary across different parts of their overlapping ranges, and in some regions, these interactions might even be absent. Despite the limitations posed by the lack of detailed information on species interactions, intraspecific variability, and spatial variation in interaction strength, our approach represents the most viable solution for addressing these research gaps. Furthermore, undocumented interactions are not included in the metaweb and, as a result, are also missing from our framework. Nevertheless, to our knowledge, this database remains the most comprehensive resource available on trophic interactions in Europe.

Finally, following Borda-de-Água et al. (2011) and Grilo et al. (2020), we used road density as a proxy to measure the impact of roads on wildlife, acknowledging that this metric may not fully capture the spatial variation in these effects. This choice assumes that road density generally correlates with traffic intensity across regions, which is reasonable for a continental-scale study. Incorporating traffic volume into the modeling framework could potentially provide a more accurate distinction between areas with higher and lower wildlife impacts. Unfortunately, a significant portion of the road network lacks standardized traffic volume data, limiting our ability to include this factor. Future research should prioritize integrating traffic volume data, to refine assessments of areas most at risk from road impacts on wildlife populations and ecological interactions. Yet, existing studies do not consistently demonstrate a clear relationship between traffic intensity or speed and roadkill rates (Denneboom et al., 2021; Grilo et al., 2015). Therefore, using road density might actually be a more reliable indicator of the potential for roads to disrupt biological communities than traffic speed or intensity.

Our findings indicate that the impacts of roads hold profound implications for ecosystem conservation. Roads are likely to prevent or hamper the spread of apex predators (Maxwell et al., 2016; Ripple et al., 2014). The negative correlation between its distribution and road density reflects this restrictive relationship. However, while road impacts, such as roadkill, may be exerting a lower role in directly depleting top-level species in their current distribution area, roads are likely to be constraining the functional connectivity among their populations. For example, a major threat to the Iberian and Eurasian lynxes’ dispersers is roadkill (KramerlJSchadt et al., 2004; Philips, 2020). On the other hand, the loss of prey due to high road densities may contribute to the failure of top-level species to persist throughout dispersal areas. Numerous basal species appear to encompass a significant portion of their ranges with road densities above their estimated threshold of vulnerability (Borda-de-Água et al., 2011; Grilo et al., 2020). Consequently, roads might be more noticeably depleting prey species, while generating less visible effects by limiting the presence of predators through disrupted interactions.

We suggest that roads are contributing to and probably accelerating the simplification and contraction of ecological networks (Felipe-Lucia et al., 2020; Heleno et al., 2020; Mestre et al., 2022). With the expected massive expansion of road networks, including the construction of 3–5 Mio km of new roads worldwide by 2050 (Meijer et al., 2018), these adverse impacts will likely extend to previously undisturbed ecosystems in developing regions, while exacerbating them in more developed countries. Therefore, immediate action is imperative to address the changes in ecosystems that may arise, namely to preserve food webs, as there is an imminent risk of pushing entire ecosystems beyond their safe limits (Steffen et al., 2015). This requires improving our understanding of the interconnections between transport networks and higher levels of ecological organization, such as ecological communities and their interactions.

This study carries significant implications for road planning and landscape management strategies, especially concerning protected areas and their connecting corridors. The focus for conserving top predators should revolve around establishing connectivity between high-quality habitat areas, strategically steering away from regions having high road density. However, their conservation must include measures to prevent the depletion of prey due to high road densities, otherwise top-level species may struggle to persist in the areas they seek to colonize. On the other hand, the impact of roads on intermediate and basal-level generalist species warrants a more cautious approach and study. These species have extensive environmental adaptability and a broad dietary spectrum, often having wide distributions throughout Europe. As previously mentioned, given that they interact with a disproportionately large set of coexisting species, they play a fundamental role in shaping ecological communities and influencing ecosystem functioning (Gaston, 2010). As so, their disappearance, namely due to high roadkill rates, may cause substantial losses in predator-prey interactions. However, the conservation of generalists, and particularly road mitigation targeting those species, is not always a priority. Moreover, while threatened or iconic species receive dedicated research and tailored mitigation measures (e.g., Ascensão et al., 2019b), more common and less appealing species may be disregarded from road management programs. But the prevalence of roadkill among common species is an alarming concern in our ecosystems (Ascensão et al., 2019a; Grilo et al., 2018). Despite their familiarity, these species often face high mortality rates that may tip local populations into extinction, causing a significant decline in their overall abundance and biomass. Such a decline is not just a threat to these species alone but also to all species they interact with, the ecosystem services and contributions from nature they provide, therefore being a pivotal factor contributing to the broader decline in biodiversity.

Overall, road and landscape management should account for the preservation of conditions for species persistence and movement for the different trophic levels. This includes habitat restoration and reducing road fragmentation by implementing tailored measures that bolster the resilience of the different trophic levels within their habitats and road proximity, such as different typologies of road passages (Clevenger et al., 2001; Lesbarrères and Fahrig, 2012). Such targeted species-specific approaches of habitat restoration and functional connectivity could play a pivotal role in mitigating the detrimental impacts of road networks on wildlife and preserving ecological network integrity.

## Supplementary Information

**TABLE S1.**
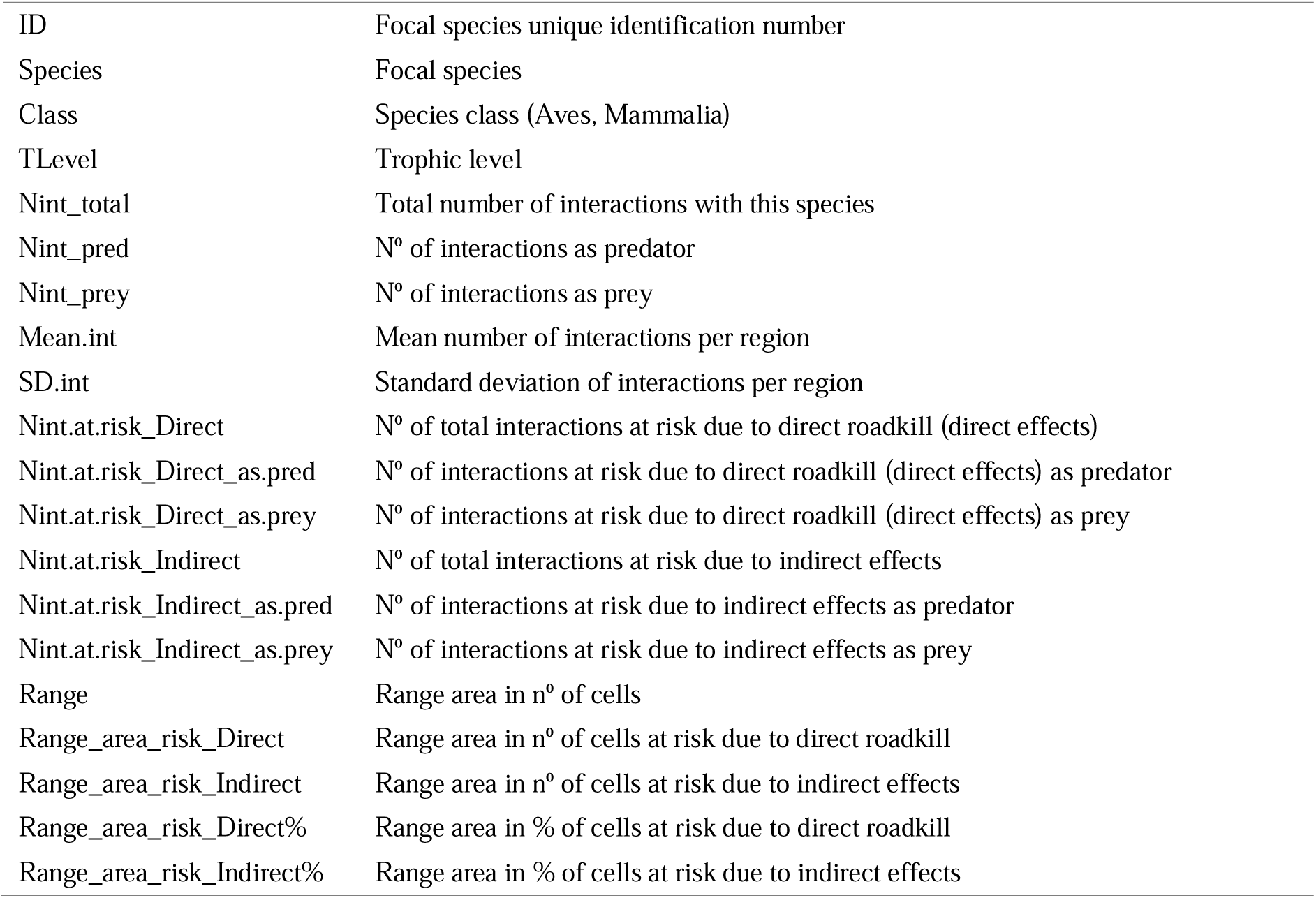
Information on species’ interaction characteristics. **Metadata of Table S1.**

## Notes

### Competing Interest Statement

The authors have declared no competing interest.

### Summary of Updates

We clarified the text, particularly the methods sections and the implications of results. Also improved figures.

